# Mechanochemical Feedback Enables Efficient Navigation in Complex Chemical Gradients

**DOI:** 10.64898/2026.07.01.735938

**Authors:** Edwin Huras, Jupiter Algorta, Henry De Belly, Orion D. Weiner, Leah Edelstein-Keshet

**Affiliations:** Mathematics Department, University of British Columbia, Vancouver, BC, Canada; Medical Center, University of Texas Southwestern, Dallas TX, USA; School of Medicine, University of California, San Francisco, CA, USA

## Abstract

Neutrophils move through narrow pores, convoluted channels, and tight spaces in tissue to find infection sites. Their ability to sense weak chemical gradients, undergo directed motion, and solve such path-finding problems rests on internal GTPase signaling circuits that control the front protrusion and rear retraction of the cell. Here we explore several variants of known core polarity circuits, with local and long-ranged negative feedback, including inhibitor downstream of Rac, Rac-Rho antagonism, and effects of membrane tension. The resulting reaction-diffusion (RD) equations for Rac and Rho are then used to simulate protrusion-retractions along the edge of a simulated motile cell. We visualize how cells navigate through narrow tracks with sharp corners and weak chemical gradients in 2D. Our metrics for cell performance include polarity initiation, wall-collision intensity, and track completion. In this way, we expose how Rac and Rho, together with their immediate down and upstream components can fine-tune neutrophil motility through complex environments.

**Author Summary:** White blood cells, attracted to sites of infection, migrate through complex tissues to find their target. Such movement requires a balance between robust polarity in one direction versus flexibility in response to spatial cues such as obstacles and sharp turns. Here we use mathematical modeling to explore known intracellular circuits that regulate front protrusion and rear retraction in directed cell migration. We test several such circuits in simulations of cells moving along zigzag tracks with sharp turns. We demonstrate that a basic cell polarity circuit, on its own, has limited success, since cells tend to get trapped in sharp corners. Known modulators of this core, which add local negative feedback, mutual front-back antagonism, and long-range feedback from membrane tension, improve cell performance. A cell with the full front-back-membrane tension regulatory circuit avoids delays due to traps and obstacle collisions, and moves swiftly through a convoluted passage to its target site.

## Introduction

Neutrophils are immune cells that respond rapidly to injury or infection, sensing subtle chemical cues and moving by chemotaxis towards the affected sites. This requires not only direction sensing, front-back polarity coordination, and motility, but also a balance between robust and flexible responses that facilitates navigation through complex tissue environments [1–4]. In this paper we ask how known intracellular regulatory components contribute to the exquisite navigation capability of such cells.

We know many of the central regulators of cell shape and movement during chemotaxis, including the Rho GTPase Rac, which modulates the protrusive front, and Rho, which mediates the contractile back, as well as phosphoinositides such as PIP3 that regulate cell polarity [6]. These circuits have complex positive and negative feedback regulation [7–11]. However, it is not clear how components of these networks synergize to solve the neutrophil path-finding problem: negotiating a way through tortuous passages, while avoiding delays at cul-de-sacs. Here we investigate known core “frontness” and “backness” intracellular circuits involved in neutrophil navigation [12, 13]. We ask how successive components contribute to fine-tuning the cell’s polarity responses to achieve effective navigation in challenging tracks with weak gradients.

To address these questions, our approach complements recent experimental work [14, 15] using mathematical modeling to demonstrate how modifications of chemotactic regulator wirings can lead to distinct emergent behaviors such as gradient sensing and navigating around mechanical barriers. By focusing on these core components, mathematical models of chemotaxis paired with experimental observations have been crucial in probing the underlying mechanisms involved in this process, often in ways that would be difficult to do with real cells [16–18]. In contrast with “top down” models in the sense of [19] (for example, [20]), we focus on a “bottom-up” approach using mechanisms known to play a role in cell polarity and motility. We start with a simple Rac “frontness” circuit [21] (depicted by an existing polarity model) and show how stepwise inclusion of known components can alleviate shortcomings, finetune the response, and allow cells to navigate complex environments. We explore next-generation models for front-back and mechanochemical communication that could enable proper cell guidance in these contexts.

Migrating cells have protrusive fronts that are organized by Rac and contractile backs that are organized by Rho. For efficient movement, the activities of these GTPases must be properly localized relative to one another and must be responsive to chemoattractants, chemorepellents, and mechanical barriers. A common framework for studying these GTPase dynamics is reaction-diffusion (RD) models, namely partial differential equations (PDEs) that describe the interactions and redistribution of molecular components [22–24]. For example, the Wave-Pinning (WP) model proposed by Mori, et al. [25] demonstrated that GTPase autoactivation, a limited total GTPase pool, and sluggish diffusion of active GTPase along the membrane suffices to explain the formation of a zone of active Rac (or Rho) at the front (or rear) of a cell; that is, such biological attributes can already account for basic cell polarization. Polarized Rac (or Rho) activity, by promoting actin assembly (through WASp family members or formins), can then generate directional movement [26–29].

The minimal WP model provides a convenient starting point for our Rac frontness circuit, even though we recently showed [30] that it falls short of explaining the full complexity of neutrophil optogenetic responses. The advantage of using WP as the starting point is that, while mimicking a basic form of polarization, it allows us to assess the roles of additional regulatory components that shape and tune cell responses. Here we seek to investigate how minimal additional effectors and feedbacks, such as the local negative feedback circuit that operates at the front [14] or the long-range mechanochemical coupling between Rac at the front and Rho at the back [31], endow the cell with finer and more sensitive responses, facilitating navigation around complex mechanical barriers.

Algorta et al. [30] compared three variants of the Rac “frontness” circuit—the base WP model, the WP with inhibitor (WPI), and the WPI-PIP3 model which includes the upstream PIP3. They simulated the three variants on the edge of motile “cellular Potts model” (CPM) cells, assumed that Rac locally promotes protrusion, and assessed the predicted responses to optogenetic stimuli of Town and Weiner [14]. In addition to fitting the three models to experimental data from [14], Algorta et al [30] demonstrated that PIP3 was required to replicate experimental results involving composite optogenetic stimuli and “exotic” cell turning behavior.

Here we extend the findings of Algorta et al [30] to probe simulated cells navigating sharp turns in response to subtle chemical gradients [32]. We start the exploration with the same Rac (“cell front”) circuits modeled by Algorta et al. [30], using their CPM formalism, available in the Morpheus software package [33]. In contrast to Algorta et al, here the model cells are subjected to subtle chemical gradients, rather than optogenetic stimuli and challenged by narrow tracks with sharp turns. This starting point provides the advantage that the model equations (PDEs) with biologically-based parameter values have already been fit, and the models validated against experimental data in [14].

Importantly, we also extend these “front” Rac motif models in two ways. First, cells do not only protrude at the front; Rho-mediated contraction at the back is important for proper neutrophil migration [31]. Therefore, we include the Rho “rear” circuit. Rho promotes myosin-mediated contraction at the rear of the cell, and is mutually antagonistic with Rac. The Rho-Rac pair are recognized as dueling competitors in many studies [12, 24, 34–37]. Furthermore, a mutually antagonistic circuit can increase the robustness of polarity in comparison to a simple positive feedback circuit such as WP [38].

Second, and more vitally, we also model the effect of membrane tension, which is known to promote Rho (at the expense of Rac) [39, 40]. As shown by de Belly et al [31], membrane tension adds rapid global feedback that has a fundamental regulatory effect: Rac activation generates protrusion, rapidly increasing membrane tension which serves as a long-range signal that promotes Rho everywhere [41, 42]. Increased Rho activity dampens Rac activation, limiting the dominance of the “front”.

More broadly, while the front (Rac-driven) and rear (Rho-driven) motifs can each contribute to directed cell motion, how these two modules coordinate to enable robust yet adaptable navigation remains an open question [11]. Such questions also naturally connect to how cells may have evolved internal control strategies to solve specific challenges posed by complex environments.

The present study addresses this gap by comparing the chemotactic success of model cells endowed with each of the three frontness circuit variants in narrow, convoluted channels. We focus specifically on evaluating the cells’ abilities to follow gradients around sharp turns and to avoid boundary collisions. We further implement a Rac-Rho mutual antagonism model with and without a membrane-tension feedback mechanism to explore the synergy between “front” and “rear” regulatory motifs. This expanded set of models enables a more comprehensive assessment of how biochemical and mechanical cues contribute to effective navigation in challenging environments.

Our analysis reveals that successful chemotaxis in complex environments depends on finding a balance between robust and flexible polarization. Although robust polarization is helpful in noisy or low-signal environments, excessive rigidity—such as “front-locking”, where cells become stuck with a fixed leading edge [14, 43], prevents effective reorientation when the gradient shifts around corners. Less successful model cells experience boundary collisions and/or trapping in corners. Across all models, we find that some form of negative feedback, whether biochemical or mechanical, is needed to destabilize the polarized front and allow for the adjustments required to successfully manage a complicated gradient-following task. By quantifying model performance using a simple yet effective collision metric, our aim is to provide a systematic evaluation of which regulatory features contribute most significantly to successful navigation in complex environments.

## Mathematical models and concepts

We first briefly review the construction of our reaction-diffusion models for Rac at the cell front, and then develop our methods and extensions.

### Cell front regulation by Rac GTPase

Three mathematical variants of the “front” (Rac) regulation (see Figure 2), based on the wave-pinning model, were implemented in the Morpheus simulation environment [33], as described in [30] and in the Supplementary Information:

**Fig 1.**
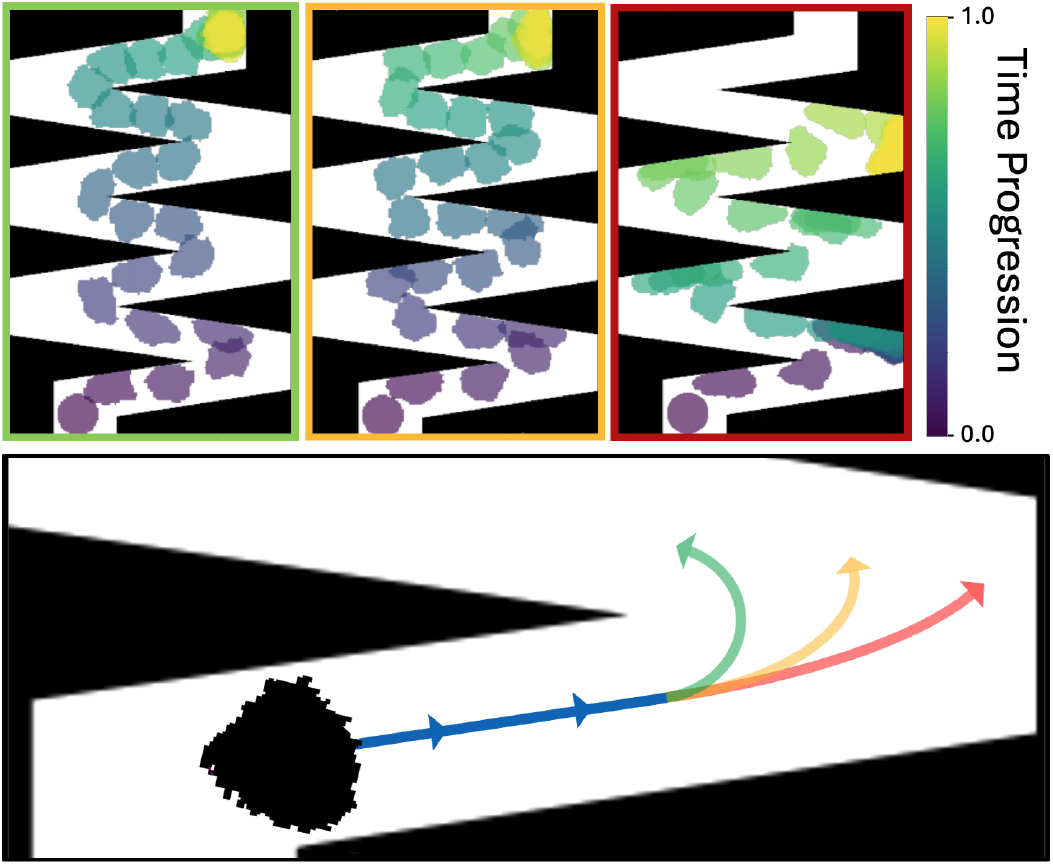
Comparing path following ability of simulated cell polarity circuits. The central question of our study is how polarity circuit variants contribute to efficient chemotaxis in confined environments with sharp turns. (See Figure 4 for the gradient.) The top panels show representative trajectories from three “frontness” Rac circuit variants with different turning abilities (Red: WP, Yellow: WPI; Green: WPI-PIP3). Each circuit is modeled as a reaction-diffusion system along the cell edge in Morpheus [33], and Rac activity promotes protrusion. Snapshots of cell positions are color-coded by time to visualize progression through the track. The bottom panel highlights the central challenge—how quickly a cell can reorient its polarity to track changes in gradient direction. Successful navigation requires both robust and flexible polarization.

**Fig 2.**
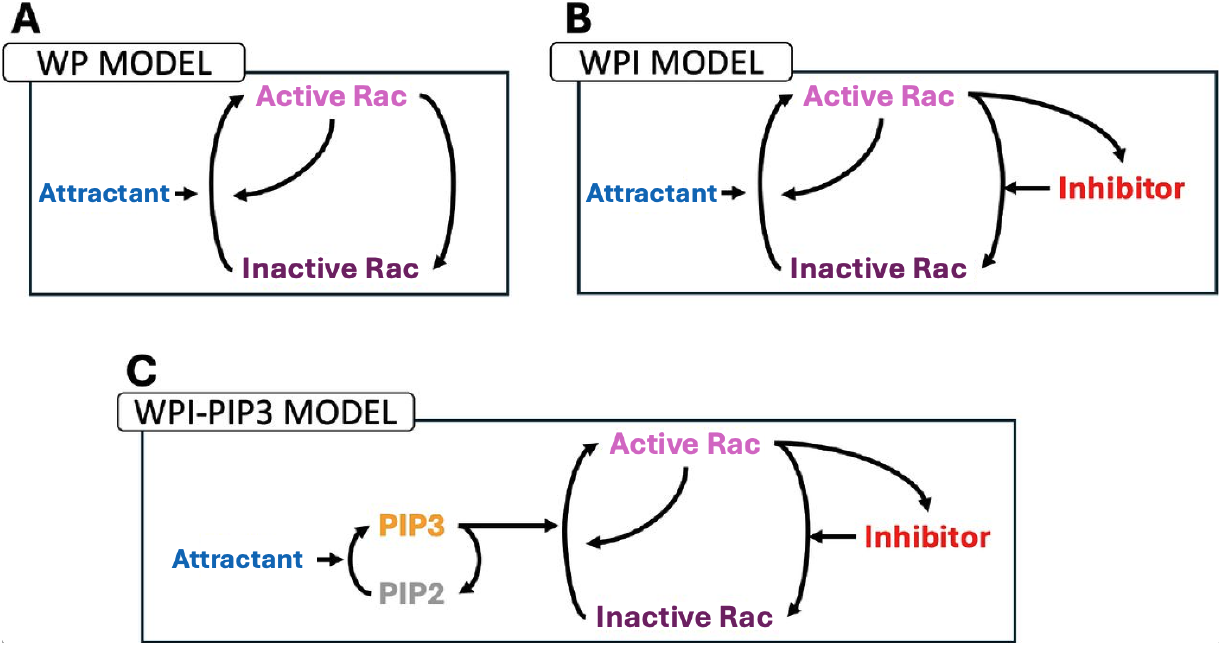
Cell front (Rac) regulatory circuits are variants of the Wave-Pinning model. Schematic diagrams for all three variants are shown. WPI includes a Rac inhibitor downstream of Rac. WPI-PIP3 further adds the PIP3 circuit. Figure modified from [30] by replacing an optogenetic signal by a chemical attractant.

#### 1. Wave Pinning (WP) model

The original mass conserved reaction-diffusion system of Mori et al. [25], shown schematically in Figure 2A. Here, a single GTPase (Rac) cycles between active and inactive forms. Active Rac, bound to the plasma membrane, diffuses slowly relative to inactive Rac. Nonlinear positive feedback (Rac self-activation), and a constant total pool of Rac are assumed. The model generates stable cell polarity by promoting a zone of high Rac activity (at the expense of the relatively uniform pool of inactive Rac). Previous work showed that this model can account for basic polarization and repolarization responses to stimuli above some threshold [25, 44]. The threshold allows for cells to have a stable “rest state” in absence of polarizing stimuli.

#### 2. Wave Pinning with Inhibitor (WPI) model

An extension of the Rac circuit that incorporates a local Rac inhibitor downstream of active Rac (Figure 2B). While its identity is as yet undetermined, the presence of such an inhibitor was demonstrated in [14] and investigated in [30], who characterized its biophysical properties through model fitting to data.

#### 3. Wave Pinning with Inhibitor and PIP3 (WPI-PIP3) model

The most detailed variant, which adds PIP3 to the WPI model, creating an additional layer of regulation (Figure C). This layer was shown (by [30] to play a role in explaining responses to composite optogenetic stimuli in [14].

Parameter values were taken from [30], who used real cell trajectory data to fit the model. Cell heterogeneity was also included, as described in the Methods. See the Supporting Information (SI) for further details on model equations, model implementation, and parameter values.

## Methods

### Multiscale (Morpheus) simulations

We used the open-source multiscale cell simulation software Morpheus [33] to simulate the RD models for regulatory components on a 1-dimensional (1D) periodic domain (mapped onto the cell edge). We also implemented a diffusion field of chemoattractant inside the zigzag channels, and simulated single cell shapes and motion through those channels using Morpheus. The zone of active Rac (Rho) at locations along the cell edge was assumed to promote front protrusion (rear retraction) where applicable. The cellular Potts model (CPM) formalism takes care of adjusting the cell rear so as to preserve the overall area of the cell near some constant “target area”. These minimal assumptions suffice to simulate cell motility. The Morpheus simulations of cell shape and motion were based on a template provided in [30]. This existing simulation environment is well suited to exploring the cellular behaviors resulting from various regulatory circuits operating along the cell edge (described by reaction-diffusion PDEs). Simulation files are available for each case.

### Domain construction and gradient establishment

To evaluate the navigation capabilities of the model variants, we constructed binary track environments in which black pixels represent impermeable regions. Using the simulation software Morpheus [33], we established chemical gradients within these tracks by placing a constant chemical source at the end point of the track and a constant sink at the entrance. For computational efficiency, we simulated a process of diffusion to steady state once, and the resulting static gradient was used from then on as a chemoattractant field for the cell to follow from entrance to end (Figure 3). This saving of computational time means that our cells do not affect the gradient.

**Fig 3.**
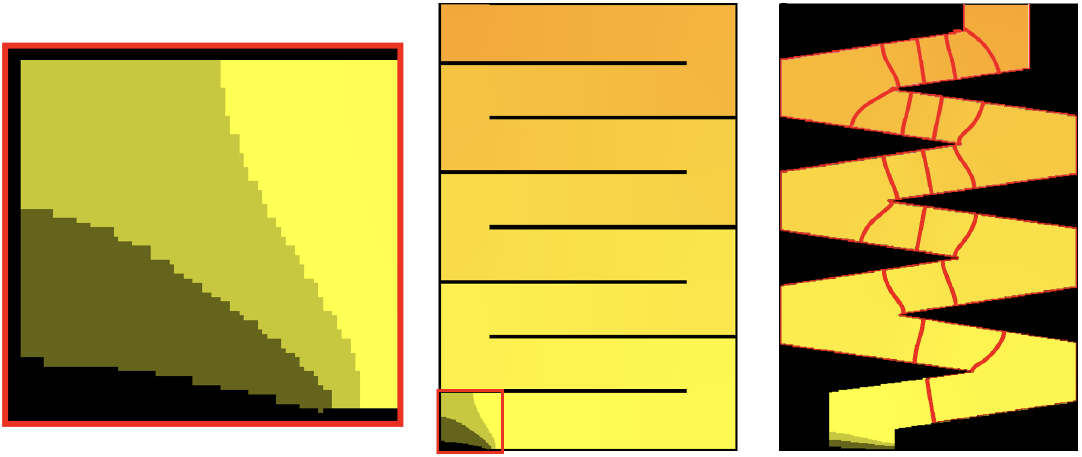
Typical chemoattractant gradient. After diffusion, the resulting gradients in the tracks were saved as 16-bit depth grayscale images that initialized the attractant gradient, with zero attractant at the start and full attractant at the end of the track. The left images represent the gradient with low concentration in yellow and high concentration in orange, and zero concentration in black to highlight the boundary. The right image highlights the staircase-like gradient cells could follow, with isolines (in red) marking loci of constant concentration at fixed increments through the track. Higher density of isolines means steeper gradients. Track B (right panel) had corners acting as traps as the gradient’s isolines become more spread out. Cells that overshoot a turn and end up in the corner often become stuck, with the external signal too weak to shift the polarization direction.

To highlight differences between the performance of the models, we developed two specialized track geometries that feature sharp turns designed to challenge the cells’ responsiveness to the gradient (Figure 4). These specialized tracks have path widths only slightly larger than the cell diameter, creating confined channels with corners that acted as traps for any cells that failed to reorient to the gradient before colliding with the boundary.

**Fig 4.**
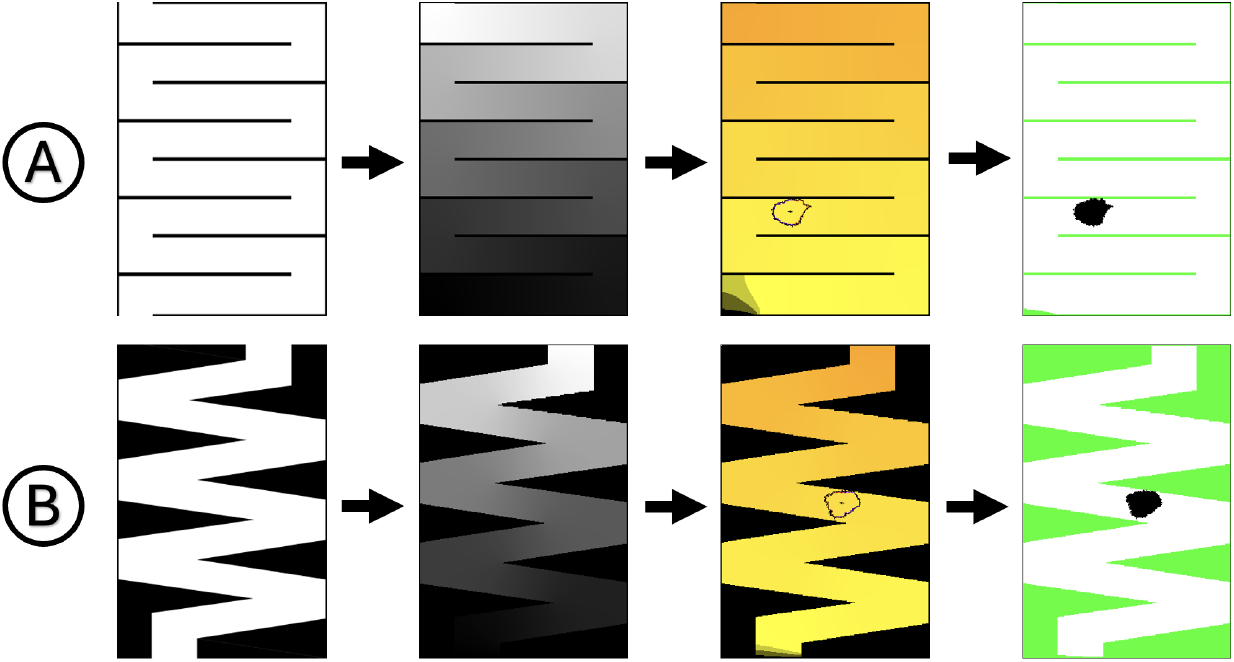
Track generation and simulation pipeline. We used two types of tracks, A (top) and B (bottom). A chemical gradient (shaded white to black and orange to yellow for high to low concentrations) was established by simulating diffusion from track exit (top) to entrance (bottom), with impermeable track walls. Chemotaxis of model cells was then simulated on each track with the fixed attractant gradient. High-contrast images of the simulated cell (black shape, right panels) were generated at fixed time steps, and then processed for cell location and collision metrics.

### Simulation procedure

We used initially uniform Rac distributions along the membrane, with small random variations to account for cell heterogeneity. Every new simulated cell would have parameter values and initial protein concentrations pulled from normal distributions with means and standard deviations taken from [30]. (see Table 1), resulting in varied behavior from cell to cell within and between models. See the Supporting Information (SI) for further details on model equations, model implementation, and parameter values.

Each model was simulated 20-30 times in each of the two specialized track geometries. For each simulation run, all important state information (cell location, protein concentrations, detected chemical signal, etc.) was recorded every 10 time-steps and high-contrast image files showing the cell and track boundaries (as in Figure 4) were saved. These data were then processed to analyze the cell’s navigation performance and boundary interactions, primarily using the image files to detect proximity to the walls and determine progress through the track.

### Collision detection and metrics

To quantify the navigation performance of each model, we developed a collision detection algorithm that identifies interactions between the cell and track boundaries by processing saved frames from each simulation. A collision was defined as occurring when any pixel of the cell’s perimeter was adjacent to a track boundary pixel. For each perimeter pixel of the cell, we examined its immediate neighborhood (the 8 pixels surrounding it) and determined whether the neighborhood intersected with the track boundary (see example in Figure 5). When an intersection occurs, the central pixel is labeled as a collision pixel.

**Fig 5.**
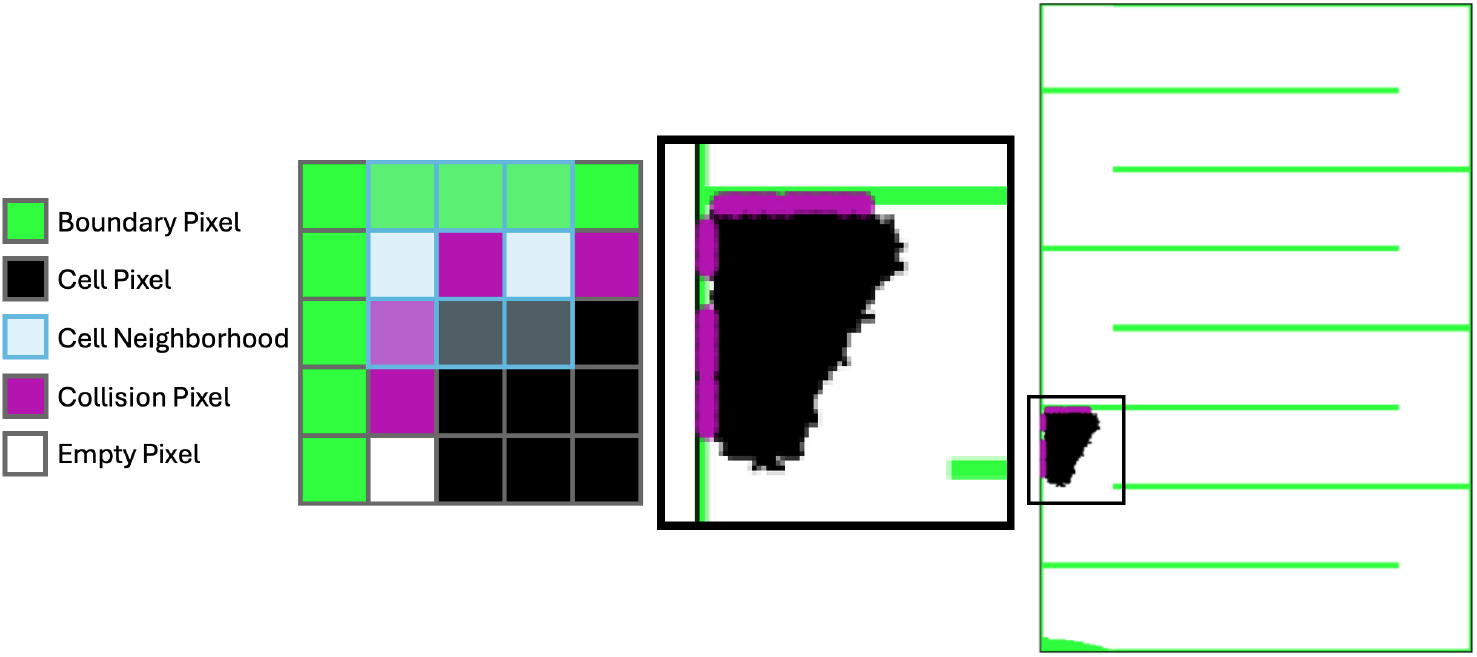
Collision intensity calculation. An illustration of collision pixels. A cell’s pixel is deemed a collision pixel (shown in magenta) if one of its 8 neighboring pixels is a boundary pixel (green). A typical cell and track image (right) and a zoomed-in view (center) are shown, enlarged for emphasis.

Using collision pixels, we then defined collision intensity, a time-dependent quantity measuring the number of collision pixels divided by the total number of cell pixels in a given frame of a simulation. The harder a cell presses into an obstacle, the more contact the cell makes with the obstacle and thus the higher the collision intensity. If a cell makes no contact with the boundary, then collision intensity is zero. While other metrics were explored (see SI), collision intensity proved to be the most sensitive at detecting when collisions occurred and to what degree the cell pressed into the obstacle.

## Results

### Qualitative observations of model behavior

We first considered several model variants for the Rac “frontness” circuit to determine whether and how the Rac downstream inhibitor and the upstream PIP3 affect the basic polarity responses captured by the Wave-Pinning model. Simulation of these three variants revealed distinct differences in the cells’ chemotactic capabilities through our track environments. Representative trajectories of each model variant can be seen in Figure 6 for Track A and Figure 7 for Track B.

**Fig 6.**
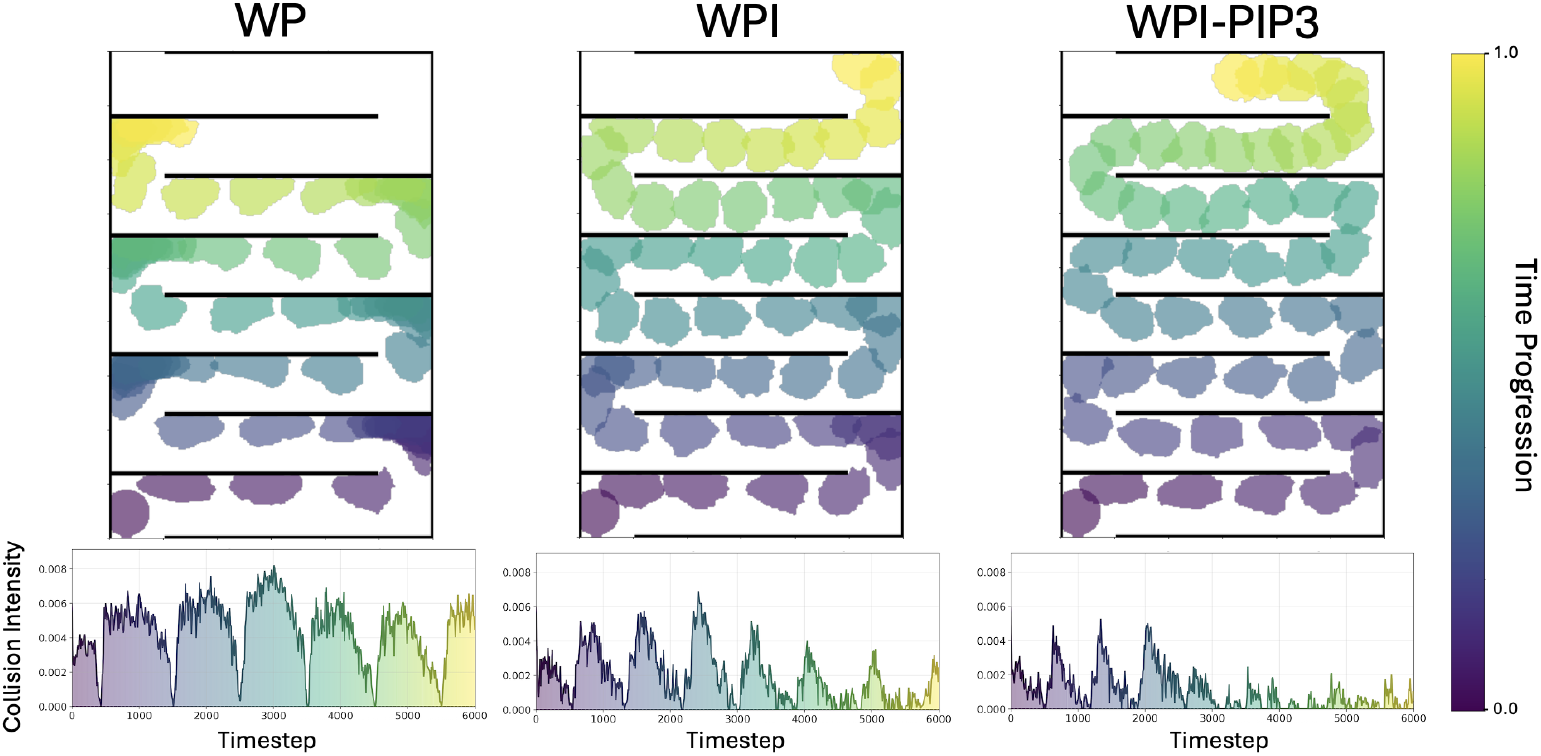
Sample trajectories of each frontness circuit and associated collision metric in Track A. The paths taken by simulated cells through the Track A geometry and associated collision intensity over time for each simulation (bottom row). Snapshots of cell position are taken every 120 timesteps. Cell snapshots and collision intensity are coloured by time progression, with all cells starting in the bottom left corner of the track. Notice the varied cell shapes and dynamics in the corners: the WP cell press into the track corners and slowly reorient while WPI-PIP3 cells mostly avoid corners by reorienting rapidly with the changing gradient.

**Fig 7.**
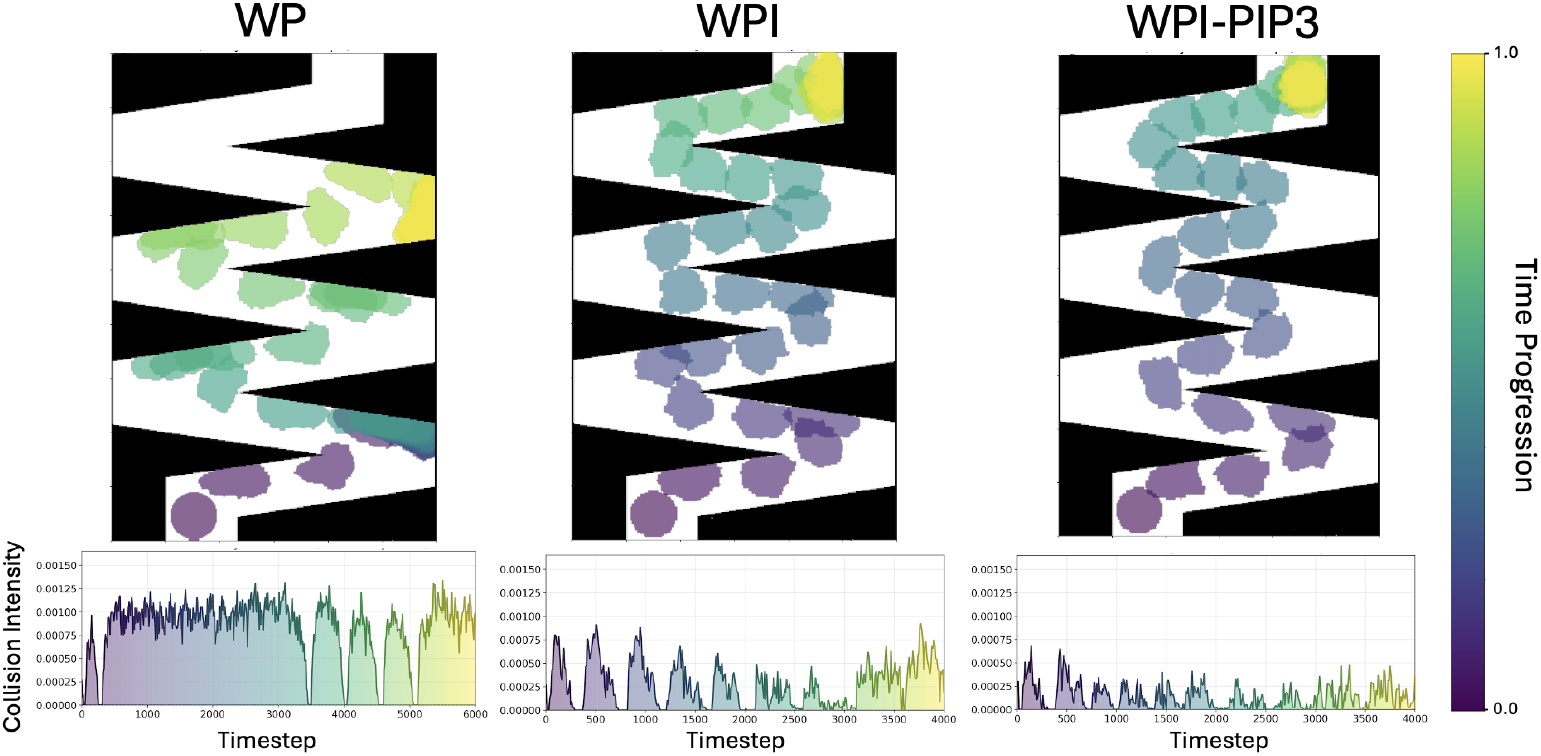
Sample trajectories of each model and associated collision metric in Track B. As in Figure 6, for track B. Note that this WP trial was an outlier, as the majority of WP cells never escaped the first corner. Clear improvement in track navigation is shown from WP to WPI and further tightening of turn radius from WPI to WPI-PIP3. Notice the extended time spent by the WP cell in the first corner, arriving around *t* = 300 and leaving around *t* = 3000. The gradient was especially subtle in the corners of Track B, trapping many WP (and some WPI) cells in the first and last corners.

Since the attractant concentration is quite weak near the starting point (nearly zero), around half of the WP model cells failed to initialize polarization at all in Track A. Those that did polarize generally exhibited strong polarization in the direction of the initial gradient but limited ability to reorient at a sharp turn. Many WP cells, overshooting the turn, got trapped in corners for extended periods of time and collided repeatedly with the track boundary. Some of these cells eventually escaped the corners by slowly reorienting in the direction of the gradient, while continuing to press into the walls. Tracking collision intensity over time for each model accurately captures this deficiency, as shown in Figure 6. We observe marked differences between the WP cell trajectories and those of the two WP variants. Collision intensity has high sustained values when the WP cell is stuck in the corners and only falls off when the cell escapes and follows the gradient down a straight section before colliding with another wall.

These results highlight the fact that a circuit composed solely of Rac and positive feedback (as represented by the traditional Wave Pinning model) is too basic to account for known neutrophil chemotactic ability. While capturing basic polarization and repolarization responses [25, 30, 44], WP lacks the robustness, adaptability, and sensitivity that the complicated path-finding task requires.

In comparison, the WPI model cell that includes a Rac inhibitor downstream of Rac shows improved reorientation capabilities. WPI cells polarized less strongly, have flatter distributions of active Rac compared to WP, and moved more slowly; interestingly, they were generally much better at navigating the sharp turns. However, WPI cells were unable to avoid the corners entirely, generally getting stuck in the first few corners before the cell slowed down in the latter half of the track. At times in Track B, cells would overshoot the turn sufficiently to get caught in a corner trap and fail to escape before the simulation timed out, similar to the end of the trajectory for WP in Figure 7.

The WPI-PIP3 model demonstrated the most effective ability to negotiate corners in both track geometries explored. WPI-PIP3 cells rarely became trapped in corners for extended periods of time, appearing to reorient rapidly when the gradient direction changes upon entering a turn. Similar to WPI, WPI-PIP3 cells slowed down further in the track, resulting in even tighter turning radii as the cell progressed.

### Rac with Inhibitor and PIP3 enables better navigation in complex environments

Across the models, the mean collision intensity for all runs varies substantially. The results are shown in Figure 8 for the two types of tracks. The WPI model shows much lower collision intensity and improved ability to negotiate a sharp corner compared to the base WP model. However, in both track geometries, the WPI-PIP3 model outperforms both WP and WPI, demonstrating significantly lower collision intensity. WPI-PIP3 model cells are more effective navigators with fewer boundary interactions. Note that only the relative collision intensity values per track are important, as the track and corner geometry influences what proportion of the cell perimeter can contact the wall.

**Fig 8.**
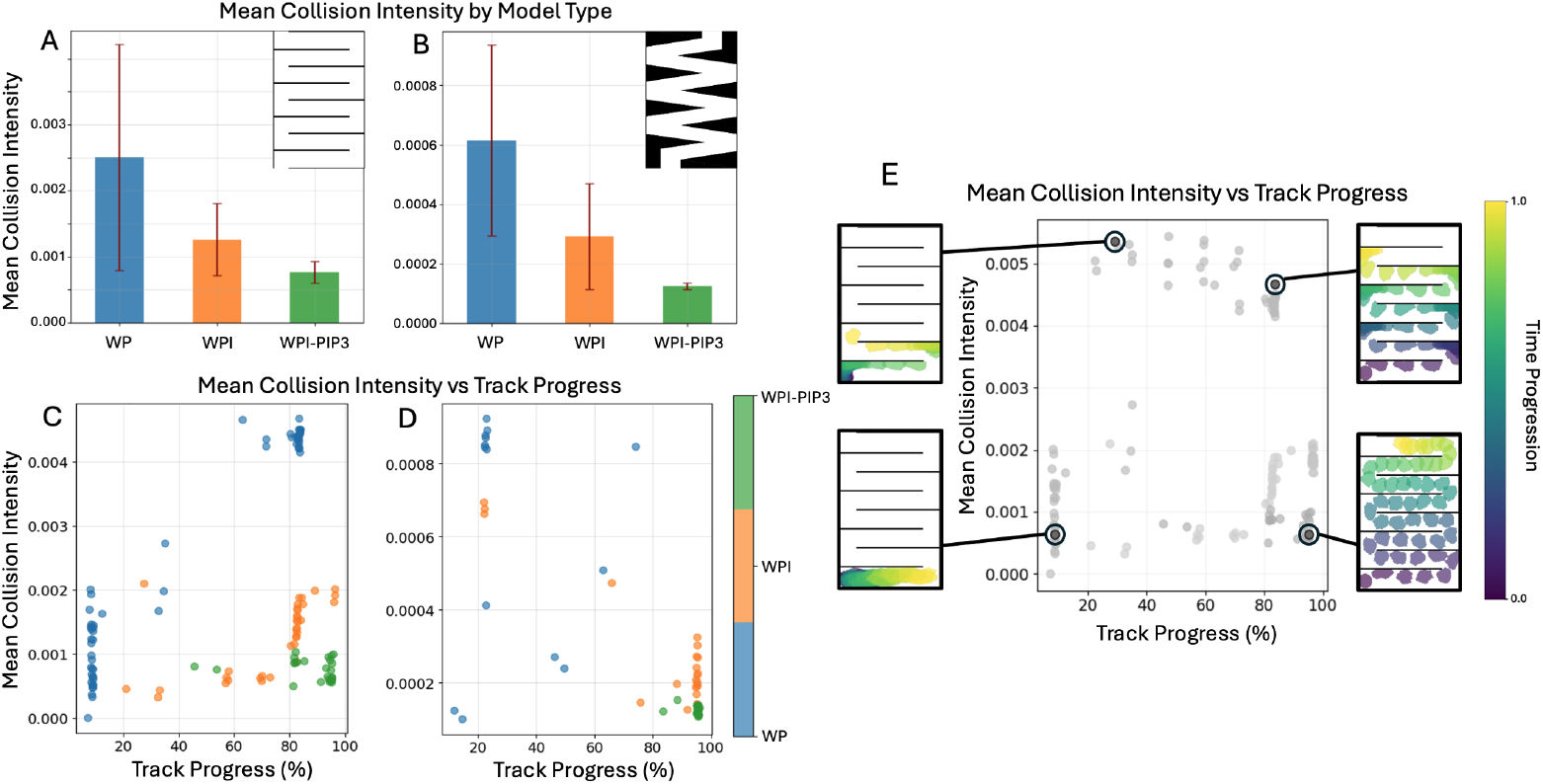
Mean collision intensity and track progress for the frontness Rac circuit variants. (**A**,**C**) Mean collision intensity by model type (WP, WPI, WPI-PIP3) for each track geometry (top right corners). Error bars indicate standard deviation across all trials. Mean collision intensity decreased and run-to-run consistency increase from WP to WPI to WPI-PIP3, consistent with improved navigation. (**C**,**D**) Individual trials use parameter values drawn from a distribution quantified from experimental data by [30] (resulting in variable runs within models to depict cell heterogeneity). Nearly half of all WP trials failed to polarize in Track A (bottom left quadrant of **C**). **(E)** Representative runs of several points on the plots. Trials in the bottom-right represent the most successful runs where cells efficiently navigated to the end of the track with limited boundary collisions. Cells that got stuck in a corner occupy the top-left region. The top-right region includes trials with frequent collisions but successful progress along the track. Cells that fail to polarize occupy the bottom-left region of the plot.

We plot individual trials by mean collision intensity versus total track progress (Figure 8), giving a better sense of how effective each individual simulated cell was at navigating the track. There is a trade-off between moving accurately (minimizing wall collisions) and moving fast enough to achieve high total track progress, with the best-performing cells capable of doing both. See Figure 8E for representative runs based on plot region.

For Track A, nearly half of the WP model cells failed to polarize sufficiently and never progressed more than 10% through the track. These cells remained mostly static, resulting in comparatively few collisions and thus relatively low mean collision intensity. The other half of WP model cells made it most of the way through the track, but with frequent and sustained track boundary collisions throughout. This explains the large variance associated with the WP mean collision intensity. In Track B, the majority of WP cells initiate movement, but proceed to get trapped in the first corner. The gradient is especially subtle in the corners of Track B, and failing to make a turn before hitting the wall generally resulted in cells getting trapped.

The WPI cells are better navigators than WP cells overall, although track progress was quite varied. Several of the lower progress trials (around 20-60% total track progress) entered into oscillatory states of stop and go motion, where the cell struggled to maintain consistent polarization, resulting in slower, pulsatile movement. These cells generally avoided track wall collisions but made little progress overall. WPI-PIP3 runs are generally consistent and always result in low collision intensity. With the exception of a few trials in Track A that entered oscillatory states similar to WPI, every WPI-PIP3 cell made it more than 80% through the tracks.

It is clear that adding the Rac inhibitor downstream of Rac significantly mitigates “front-locking” behavior in the cells, without negatively impacting their ability to polarize. The inhibitor helps destabilize the active Rac zone pinned at the front of the cell, prompting it to reorient more quickly when the external gradient starts to point in a new direction. Evidence of WPI-PIP3’s rapid turning ability was already seen in [30]. We also recall that the fine-tuning role of the phosphoinositides was already strongly emphasized by [45], though in the context of a more detailed simulation model. Our results so far suggest that building back up some of the known regulatory layers found in real cells enables us to capture behaviors that are closer to physiology.

### Front-rear circuit and the role of membrane tension

So far, we have considered only the regulation of a cell’s front by Rac and its associated proximal effectors. We next sought to add two important missing features: 1) feedback from the rear polarity circuit (whose central regulator is Rho) and (2) feedback from mechanics in the form of membrane tension. We have seen that in the absence of such influences, our simulated cells have limited feedback from the outside environment—besides the chemical gradient—and this is evident when a cell pushes persistently into a barrier. While these cells are blocked by track boundaries, the effective mechanical influence and consequent cell deformation have no impact on the internal signaling dynamics. The result is that a cell, once polarized, will continue to push in the same direction until the gradient causes it to reorient, regardless of whether that cell is making progress or stuck in a corner. The failure of the WP model to reorient shows that robust polarization by the front circuit alone does not suffice for successful navigation in a complex environment without additional feedback that can improve sensitivity.

The WP model in its simplest form includes a proto-global feedback mechanism (rapid diffusion of inactive Rac encodes global information about the cell’s overall activation state). However, clearly, the basic WP model as such was not formulated to describe a cell’s response to mechanical stimuli (but see [46, 47] for variants that do so). Simply put, direct feedback from mechanics to the signaling system is missing in our implementation of the WP model and its two variants (WPI, WPI-PIP3) so far. Importantly, [31] suggest that membrane tension provides vital rapid long-range communication across the entire cell membrane, conveying information about protrusive Rac activity at the front to the contractile Rho activity at the back. Intuitively,

Rac-based protrusion increases membrane tension globally and induces Rho activity, which temporarily damps the Rac polarization zone. We sought to explore their findings within the context of the same modeling framework of simulated motile cells.

Accordingly, we explored two front-rear (Rac-Rho) circuit variants, as shown in the schematic diagrams of Figure 9. As before, each one was represented by a set of reaction-diffusion equations along the cell edge, together with a proxy for membrane tension based on cell perimeter. The variants included

**Fig 9.**
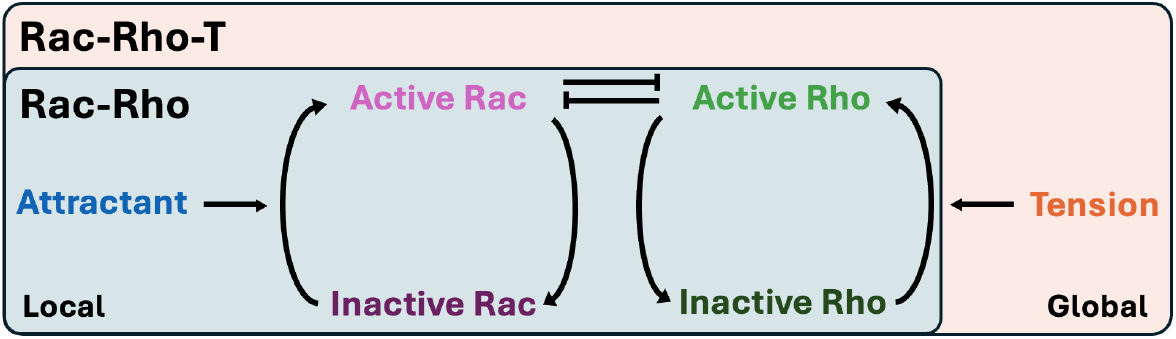
Rac-Rho model variants. Schematic diagrams for local Rac-Rho mutual inhibition models with (full box) and without (blue box) the global feedback from membrane tension. Full diagram, including global Tension, comprises the Rac-Rho-T model. Similar to the WP models, the external attractant promotes the activation of Rac. The addition of Rho creates a new dynamic, both as an antagonist to Rac and as a promoter of contraction where Rho activity is high. The Rac-Rho-T model adds a tension mechanism that, when triggered, rapidly produces active Rho globally, potentially triggering tension-based depolarization.

1. **Rac-Rho**: Rac-Rho mutual antagonism model with active and inactive forms of both Rac and Rho, coupled together via a “toggle switch” [48]. The rate of activation of each GTPase was assumed to be depressed by the active form of its antagonist. In the cell motility simulations, we assumed that active Rac at a given point on the cell edge promotes protrusion, as before, while active Rho promotes contraction. To facilitate comparison with the previous WP models, we use similar parameter distributions for the Rac-Rho RD models where possible.
2. **Rac-Rho-T**: As in the Rac-Rho model, but with feedback from membrane tension (using an increase in cell perimeter as a proxy) that globally enhances Rho activation [15, 40, 49]. Due to mutual antagonism between Rac and Rho, this tension-induced Rho activation necessarily produces global damping of Rac activity. Hence, sufficient tension can depolarize a cell.

Representative trajectories for both Rac-Rho and Rac-Rho-T are compared side-by-side along with a WP trajectory on Track A in Figure 10 and on Track B in Figure 11. The Rac-Rho model behaves very similarly to the WP model, with a few notable differences. For one, despite all cells being initialized with similar noisy uniform Rac distributions, no Rac-Rho cell failed to initiate polarization in Track A, whereas nearly half of WP models did. This finding is consistent with previous work showing that Rac-Rho (or similar mutual antagonists) results in more robust polarization than simple positive feedback [38, 50, 51]. Here we found that our Rac-Rho model cell could sense the gradient and initiate the formation of separate and opposing Rac and Rho peaks on either side of the cell. Second, the added contraction due to active Rho at the rear of the cell increases the pressure that a cell could sustain against a wall, increasing overall collision intensity whenever the cell overshot a turn.

**Fig 10.**
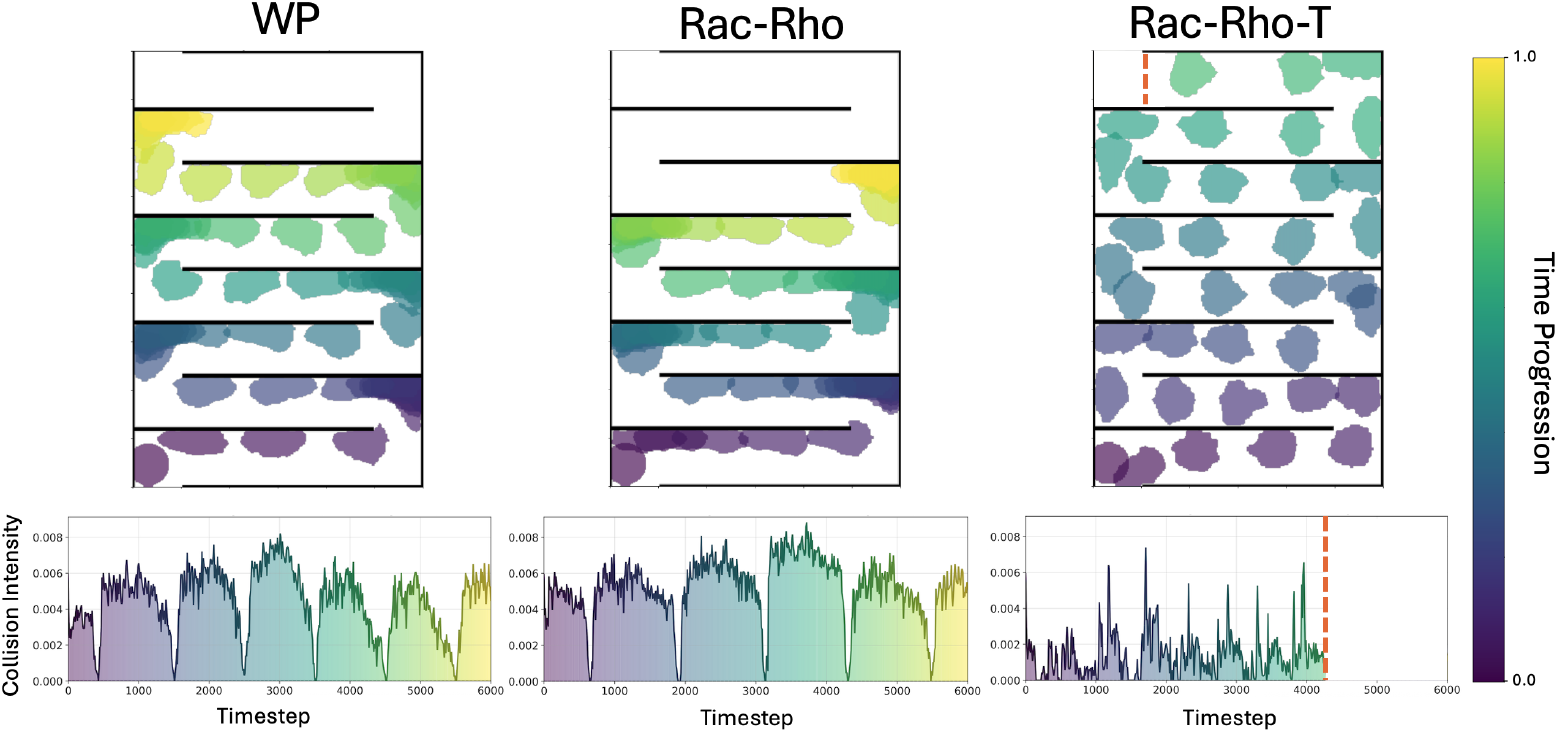
Representative trajectories for Rac-Rho front-back circuits in Track. **A**.Rac-Rho cells press more persistently into the corners due to the added Rho-dependent contraction at the rear of the cell, resulting in higher collision intensity compared to WP. Including tension in the Rac-Rho-T model variant eliminates this problem; collision with a wall causes the cell to contract due to tension-based Rho activity, reducing overall collision intensity. The orange dotted line in the Rac-Rho-T run marks the point where the cell reached the end of the track.

**Fig 11.**
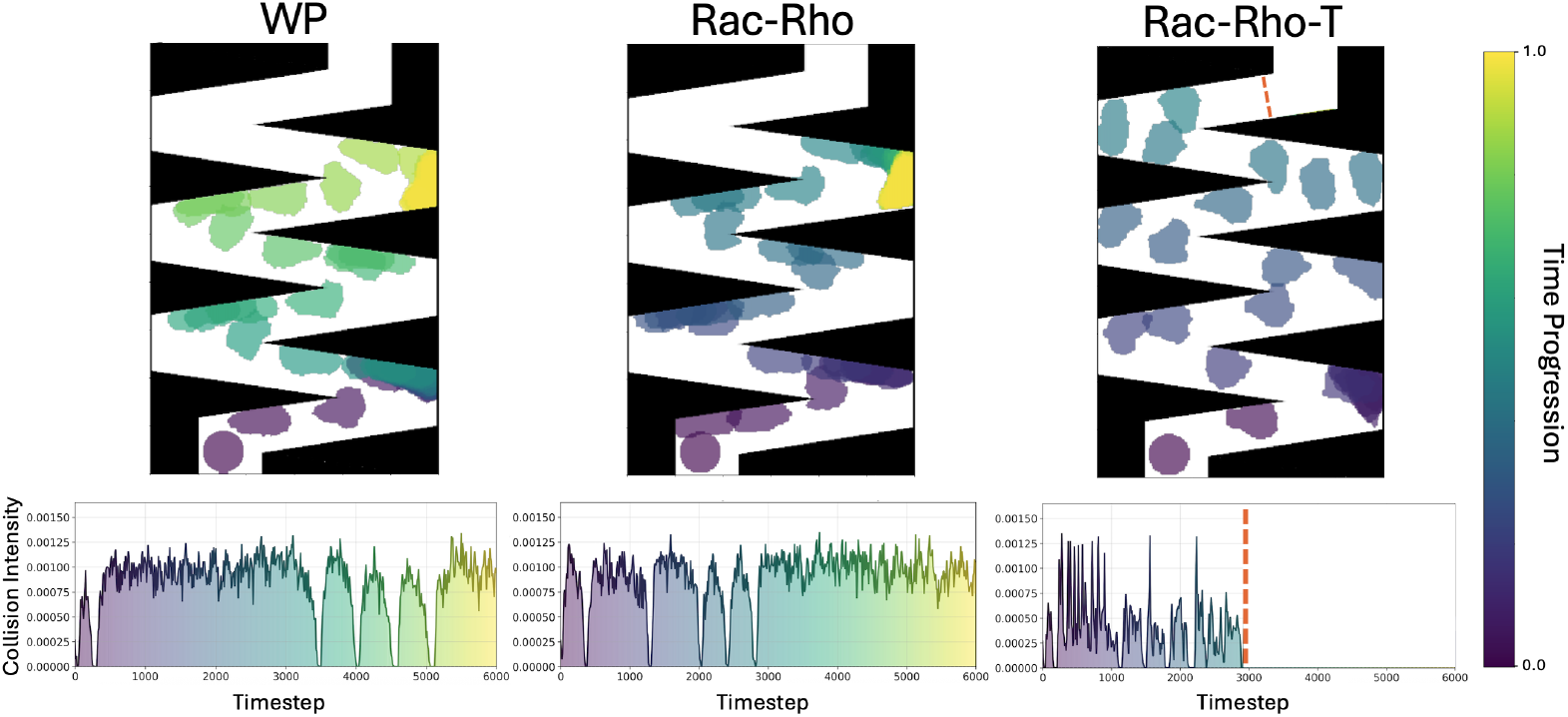
Representative trajectories for Rac-Rho front-back circuits in Track. **B**.As in Figure 10 but for Track B, with WP for comparison. Rac-Rho cells escaped the first corner more quickly, before becoming trapped in the second-last turn. Note that the WP trial shown here is an outlier, with the vast majority of runs failing to escape the first corner (see Figure 14 for individual trials). Rac-Rho-T model cells sometimes enter corner traps but succeed to escape by a sequence of tension-based depolarization events; all such cells arrived at the end of the track, here marked with the orange dashed line.

**Fig 12.**
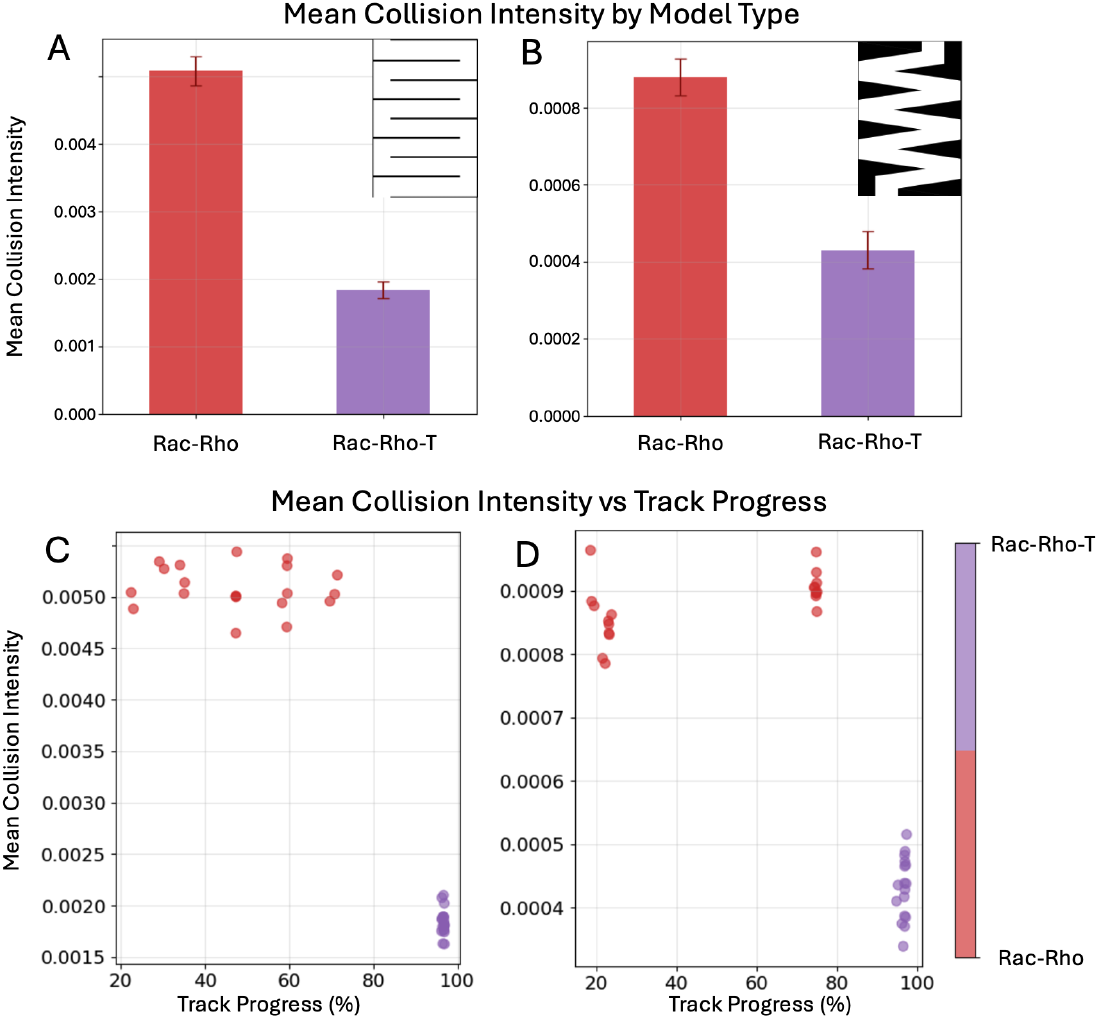
Summary results for Rac-Rho variants. (**A**,**C**) Mean collision intensity for Rac-Rho variants, for Track A (left) and Track B (right). Bars represent mean values and error bars indicate standard deviation across all trials. Rac-Rho shows very high collision intensity on average, likely due to more robust polarization and the added contraction mechanism from Rho allows the cell to press more persistently into the boundary, further increasing collision intensity. Inclusion of mechanical feedback from membrane tension results in a significant drop in collision intensity for both tracks. **(C**,**D)** Individual trials showing mean collision intensity and total track progress. Rac-Rho shows varied success in Track A, often getting stuck in corners. In Track B, there is a bimodal distribution for Rac-Rho, with cells either getting stuck in the first or second-to-last corners. In contrast, tension-mediated feedback provides a mechanism for escaping corners after collisions, a clear advantage for these cells as all Rac-Rho-T trials reached the end of both tracks.

Interestingly, the simple addition of tension leads to drastically improved results. For Rac-Rho-T model cells, overshooting a turn would result in a wall collision, just as for Rac-Rho. However, unlike Rac-Rho, these collisions do not produce sustained boundary contact, as the resulting tension spike causes the cell to temporarily depolarize and rebound from the wall.

## Discussion

Others have simulated the chemotactic motion of cells through complex tracks and mazes [52–55]. In contrast with these studies, we set out to dissect well-known minimal front and back polarity circuits of motile cells. We focused on how individual components of those circuits endow a cell with the flexibility needed to navigate complex, confined environments with sharp turns and subtle gradients. To do so, we adapted several data-based reaction-diffusion models for Rac and Rho, incorporating core signaling components and their proximal effectors We tested how these elements, alone or in tandem, shape navigation when cells respond to both chemical cues and mechanically induced feedback. Specifically, we simulated three “cell front” Rac circuits (WP, WPI, WPI-PIP3), and two “front-rear” Rac–Rho circuits (Rac-Rho, Rac-Rho-T) with and without tension-based mechanical feedback. This framework allowed us to assess how interactions among these components shape cell responses—supporting gradient detection, robust polarization, flexible reorientation to changing cues, and collision detection and avoidance.

Our results reveal a consistent theme: efficient navigation needs more than just basic polarization; it also requires the capacity for flexible reorientation when the external environment changes. The “front-locking” behavior [14, 43] observed in the basic WP and Rac-Rho models, where the cell remains persistently polarized in the original direction and fails to reorient at turns, leads to frequent trapping in corners and persistent boundary collisions that slow or trap the cell. Rather than being merely a shortcoming, these failures are informative: they suggest that a minimal, proto-polarization module (WP) could support biased movement toward attractive cues but lacks robustness and tends to get stuck in more complicated environments. Adding a Rac-linked inhibitor can be viewed as a step toward fine-tuning front regulation to increase flexibility, with additional lipid feedback (e.g., PIP3) further improving reorientation. Incorporating a rear (Rho) module strengthens polarity robustness— sometimes at the cost of ineffective pushing into corners—while membrane-tension feedback helps coordinate front–rear dynamics to rescue this synergy and produce the most effective navigators.

Several simplifications in our study merit mention. Our models represent cells in two dimensions and solve reaction-diffusion equations only along a 1D circular domain, projected onto the cell edge. Clearly, our models abstract and simplify internal cytoskeletal reorganization (assuming merely Rac and Rho dependent forces of protrusion and retraction). We also stripped away other potentially important aspects of cell biology to focus on core signaling components. The biophysical details of membrane deformation, curvature-dependent protein signaling, and details of mechanical coupling are also absent, as is the full complexity of real signaling networks present in migrating cells. However, by using relatively simple mathematical models and focusing on core features, we gain more insight into what properties and mechanisms are necessary for basic cell navigation in complex confined channels. Our results should hopefully inspire new live-cell experiments. In particular, it would be useful to study how real cells respond to similarly convoluted tracks, and how application of various inhibitors of Rho or membrane tension or Rac/Rho crosstalk and positive/negative feedback affect the successful chemotaxis of the cells in such challenging environments.

## Acknowledgments

This work was supported by a Natural Sciences and Engineering Research Council (NSERC, Canada) Discovery Grant (LEK), an NSERC Undergraduate Student Research Award (EH), and National Institutes of Health grants R35GM118167 (ODW) and R00GM154115 (HDB).

