## Supporting Information for "Mechanochemical Feedback Enables Efficient Navigation in Complex Chemical Gradients"

### Mathematical Models

#### Reaction-Diffusion Equations

The models in our paper are reaction-diffusion equations, implemented on a one-dimensional (1D) periodic ("wrap-around") domain ($\Omega$) representing the cell edge. The "frontness" Rac models were previously described by Algorta et al [30], and we briefly summarize those below. The Morpheus implementation of these models is explained further on.

#### Wave-Pinning (WP) Model

This model was originally derived to describe cell polarization based on a single GTPase [25], here we model Rac a type of Rho GTPase. Let $u(x,t)$ and $v(x,t)$ denote the levels of active and inactive Rac at position $x$ along the cell edge at time $t$. A term, $S(x,t)$, is added to the rate of Rac activation to account for an external stimulus [44]. While in [30] the stimulus was optogenetic, here, $S(x,t)$ represents the level of attractant sensed at a given location along the cell membrane, mapped from the adjacent value of an extracellular chemical field. The governing equations for Rac in the WP model are:

$$\frac{\partial u}{\partial t} = v\left(k_0 + \alpha_S S(x,t) + \gamma \frac{u^n}{K^n + u^n}\right) - \delta u + D_u \frac{\partial^2 u}{\partial x^2}, \tag{1a}$$

$$\frac{\partial v}{\partial t} = -v\left(k_0 + \alpha_S S(x,t) + \gamma \frac{u^n}{K^n + u^n}\right) + \delta u + D_v \frac{\partial^2 v}{\partial x^2}, \tag{1b}$$

where the parameters are $k_0$ (basal Rac activation rate), $\gamma$ (maximal Rac autoactivation rate), $K$ (Rac level inducing 50% maximal autoactivation), $n$ (Hill coefficient, $n = 2$), $\delta$ (basal Rac inactivation rate), $D_u$ and $D_v$ (diffusion rates of active and inactive Rac, respectively). Default values of these parameters are given in Table 1.

Equations (1) have the property that the total amount of Rac, $T$, is constant in the domain

$$T = \int_\Omega (u+v)dx. \tag{2}$$

The parameter $T$ is (indirectly) set by picking initial distributions for $u, v$ at time $t = 0$. Further information about the WP system can be found in [25,30,56].

The RD dynamics are implemented in one spatial dimension (1D), as Morpheus does not currently support RD solvers on deforming 2D or 3D domains. This 1D simplification is similar to prior works by [43] and [57]. However, in practice, Morpheus solves the RD equations on the perimeter of a circle and maps those solutions onto the deforming cell edge.

#### Wave-Pinning with Inhibitor (WPI) Model

The Rac inhibitor, $h(x,t)$ was proposed by Town and Weiner [14] and modeled by [30]. For simplicity, Algorta et al [30] assumed that the inhibitor is produced at a rate proportional to active Rac ($u(x,t)$), and degrades with first-order kinetics, leading to the model equation

$$\frac{\partial h}{\partial t} = k_h u - \delta_h h + D_h \frac{\partial^2 h}{\partial x^2}, \tag{3a}$$

where the inhibitor parameters are: $k_h$ (inhibitor production rate), $\delta_h$ (degradation rate), and $D_h$ (inhibitor diffusion coefficient). The extended model includes a modification of (1), assuming that Rac inactivation increases linearly with the level of the inhibitor, $h$. The new equations for the WPI version of the model become:

$$\frac{\partial u}{\partial t} = v\left(k_0 + \alpha_S S(x,t) + \gamma \frac{u^n}{K^n + u^n}\right) - (\delta + k_1 h)\, u + D_u \frac{\partial^2 u}{\partial x^2}, \tag{3b}$$

$$\frac{\partial v}{\partial t} = -v\left(k_0 + \alpha_S S(x,t) + \gamma \frac{u^n}{K^n + u^n}\right) + (\delta + k_1 h)\, u + D_v \frac{\partial^2 v}{\partial x^2}. \tag{3c}$$

Here, the inhibitory term $k_1 h$ introduces a negative feedback loop, with $k_1$ controlling the strength of inhibition. Algorta et al [30] showed that the WPI model successfully captures adaptive features observed in experiments.

#### Wave-Pinning with Inhibitor and PIP3 (WPI-PIP3) Model

To explain neutrophil responses to composite optogenetic stimuli [14], Algorta et al [30] showed that the upstream effect of PIP3 had to be included. This led to inclusion of this additional variable.

Denoting the level of PIP3 by $p(x,t)$, and assuming it is produced from a basal pool of PIP2 in response to an external stimulus leads to the equation

$$\frac{\partial p}{\partial t} = (T_p - p)(k_p + \beta S(x,t)) - \delta_p p + D_p \frac{\partial^2 p}{\partial x^2}, \tag{4a}$$

where the model parameters are: $T_p$ (total local pool of PIP3 and PIP2), $k_p$ (basal PIP3 production rate), $\beta$ (stimulus-enhanced rate of PIP3 production), $\delta_p$ (basal PIP3 decay rate), and $D_p$ (PIP3 diffusion coefficient). This equation assumes a constant level $T_p$ of PIP2 and PIP3 available everywhere, rather than a globally conserved pool. This is a simplification, as modeled by Algorta et al. [30], that neglects spatial variation in PIP2. Taking PIP3 into account modifies the Rac model, leading to the new system of equations (3a), (4a), and

$$\frac{\partial u}{\partial t} = v\left(k_0 + \alpha_p p(x,t) + \gamma \frac{u^n}{K^n + u^n}\right) - (\delta + k_1 h) u + D_u \frac{\partial^2 u}{\partial x^2}, \tag{4b}$$

$$\frac{\partial v}{\partial t} = -v\left(k_0 + \alpha_p p(x,t) + \gamma \frac{u^n}{K^n + u^n}\right) + (\delta + k_1 h) u + D_v \frac{\partial^2 v}{\partial x^2}. \tag{4c}$$

This system forms the WPI-PIP3 model. The key idea is that PIP3 introduces a time delay between external stimulation and Rac activation, spreading the effect of the attractant signal over time and spatially across the cell membrane. See Algorta et al. [30] for match with experimental results on cell trajectories in local-then-global optogenetic stimulation protocols.

#### Rac-Rho Mutual Antagonism Model (Rac-Rho)

The Rac-Rho models discussed in our paper are related to those studied theoretically in [50]. De Belly et al. [31] explored a mechanochemical, mutually antagonistic Rac-Rho PDE model, with active and inactive forms of both Rac and Rho. We here implement a variant of their Rac-Rho model. The primary difference is how the external stimulus affects the equations. Note that for our implementation, the external stimulus $S(x,t)$ (the chemoattractant concentration at a location along the cell membrane) only directly promotes conversion of inactive Rac to active Rac. The attractant has no direct effect on Rho. We also modified the parameters to better align with our existing model variants.

$$\frac{\partial R}{\partial t} = R_i\left(k_0 + \alpha_S S(x,t) + \gamma \frac{K^n}{K^n + \rho^n}\right) - \delta R + D_R \frac{\partial^2 R}{\partial x^2}, \tag{5a}$$

$$\frac{\partial R_i}{\partial t} = -R_i\left(k_0 + \alpha_S S(x,t) + \gamma\frac{K^n}{K^n+\rho^n}\right) + \delta R + D_{R_i}\frac{\partial^2 R_i}{\partial x^2}, \tag{5b}$$

$$\frac{\partial \rho}{\partial t} = \rho_i\left(k_{0,\rho} + \gamma_\rho\frac{K^n}{K^n+R^n}\right) - \delta_\rho\rho + D_\rho\frac{\partial^2 \rho}{\partial x^2}, \tag{5c}$$

$$\frac{\partial \rho_i}{\partial t} = -\rho_i\left(k_{0,\rho} + \gamma_\rho\frac{K^n}{K^n+R^n}\right) + \delta_\rho\rho + D_{\rho_i}\frac{\partial^2 \rho_i}{\partial x^2}. \tag{5d}$$

The parameter $K$ represents the activation threshold for the mutual inhibition term. Because this parameter was not fitted to empirical data, we assumed $K = 1$ for simplicity, reducing the Hill-type terms to the form $\frac{1}{1+\rho^n}$ and $\frac{1}{1+R^n}$.

Previous work has shown that Rac-Rho (or similar mutual antagonists) results in more robust polarization than simple positive feedback [24,38,51].

#### Rac-Rho Mutual Antagonism Model with Tension (Rac-Rho-T)

This variant builds on the Rac-Rho model by including feedback from membrane tension that globally enhances Rho activation. Only the PDE for Rho is modified, to the new version,

$$\frac{\partial \rho}{\partial t} = \rho_i\left(k_{0,\rho} + \gamma_\rho\frac{K^n}{K^n+R^n} + \frac{\beta\,\text{Tension}^{n_T}}{T_0^{n_T}+\text{Tension}^{n_T}}\right) - \delta_\rho\rho + D_\rho\frac{\partial^2 \rho}{\partial x^2} \tag{6a}$$

where $\beta, T_0$, and $n_T$ are parameters that tune the effect.

As shown by Magno et al. [58], the actual physical tension $T_{\text{phys}}$ exerted by the cell membrane in a Cellular Potts Model is derived from the perimeter constraint in the Hamiltonian as $T_{\text{phys}} = 2\lambda_p(\text{Perimeter} - \text{Target Perimeter})$, where $\lambda_p$ is the strength of the constraint. Since we are not primarily concerned with the exact details of forces, and $\lambda_p$ is kept the same across all our simulated cells, we model the mechanochemical feedback by assuming the tension signal driving Rho activation is proportional to this physical tension. Specifically, we use the raw perimeter extension:

$$\text{Tension} = \max(0, \text{Perimeter} - \text{Target Perimeter}). \tag{6b}$$

By using this proportional measure as the input to the Hill function, the magnitude of the mechanochemical feedback can be tuned via the single parameter $\beta$. Due to mutual antagonism between Rac and Rho, this tension-induced Rho activation necessarily produces global damping of Rac activity. Hence, sufficient tension can depolarize a cell.

### Model Implementation

#### The Cellular Potts Model (CPM) Simulations

To simulate cell motility driven by Rac dynamics, we implemented the model equations (1) on the edge of a 2D deformable simulated cell in Morpheus [33]. Morpheus is an open source multiscale modelling platform for simulating single and collective cell behavior, based on the Cellular Potts Model (CPM). For fuller details about the Cellular Potts Model, see [59].

The advantage of Morpheus is that most of the technical details are handled in the background by the software and plugins. Hence, the summary below is only provided as background reference.

CPM cells are represented by a connected set of pixels with a fluctuating edge that can randomly protrude or retract at each point. An energetic cost function (the "Hamiltonian") is defined as

$$H = \lambda_a(A - A_0)^2 + \lambda_p(P - P_0)^2.$$

where $A_0, P_0$ are target area and perimeter and $\lambda_a, \lambda_p$ are strengths of the area and perimeter constraints.

In our simulations, we set $\lambda_a = 1$ and $\lambda_p = 0.8$. We also set the asphericity $\phi = 1.2$. These parameter values are similar to those used by Algorta et al. [30] ($\lambda_p = 1$, $\phi \approx 1.4 - 1.46$). Aspherity is defined as

$$\text{Aspherity} = \phi = \frac{P_0}{2\sqrt{\pi A_0}}.$$

The cellular Potts simulation chooses random pixels at the cell edge and attempts swaps ("spin flips") between intra and extracellular pixels. A biasing energy term $H_b$ is incorporated in the decision to accept or reject a pixel flip.

In the cell motility simulations, we assumed that active Rac locally promotes protrusion, while active Rho promotes contraction. The probability of accepting a protrusion attempt is

$$P(\Delta H, H_b) = \begin{cases} P_{\text{protrude}} = 1, & \text{if } (\Delta H - H_b) \leq 0, \\ P_{\text{protrude}} = \exp\left(-\frac{(\Delta H - H_b)}{T}\right), & \text{if } (\Delta H - H_b) \geq 0, \end{cases} \tag{7}$$

where $T$ is a parameter that controls random fluctuations (analogous to "temperature" in physics).

The probability of accepting a retraction attempt is

$$P(\Delta H, H_b) = \begin{cases} P_{\text{retract}} = 1 & \text{if } (\Delta H + H_b) \leq 0, \\ P_{\text{retract}} = \exp\left(-\frac{(\Delta H + H_b)}{T}\right), & \text{if } (\Delta H + H_b) > 0. \end{cases} \tag{8}$$

In our simulations, we define a single net outward bias field $H_b$ along the cell edge based on the local concentrations of active Rac ($R$) and active Rho ($\rho$). Because active Rac promotes protrusion and active Rho promotes contraction, the net outward bias at a given edge location is defined as:

$$H_b = \lambda(R - \rho) \tag{9}$$

where $\lambda$ is the strength of the bias (implemented via the StarConvex plugin in Morpheus).

Under this formulation, at a location with high Rac activity, the outward bias $H_b$ is positive. This decreases the effective energy barrier $(\Delta H - H_b)$, making an outward protrusion more likely to be accepted. Conversely, at a location with high Rho activity, the outward bias $H_b$ becomes negative. In the case of an attempted retraction, the effective energy change is $(\Delta H + H_b)$; adding the negative $H_b$ lowers the energy barrier, making retraction more likely.

### Morpheus implementation

Morpheus cannot currently solve RD equations on deforming 2D or 3D domains as formerly done in [45, 60, 61]. The Morpheus plugin MembraneProperty enables a system of reaction-diffusion dynamics on a circular domain with the same area as the CPM cell that is then "projected" onto the deforming cell edge. The Morpheus plugin StarConvex was used to favour protrusion (retraction) at edge locations with high Rac (Rho). Note that in recent versions of Morpheus, the StarConvex plugin has been removed and replaced by the SurfaceMotion plugin. We also used the Morpheus Protrusion plugin. This plugin implements a persistence in locomotion based on the actin-turnover delay ("Act") idea of Niculescu et al. [62]. The RD system is implemented in Morpheus using the DiffEqn plugin, which allows direct numerical integration of the model PDEs.

We set the Monte Carlo Step (MCS) duration to 0.3 and the system time-scaling to 5 for the main RD systems (except the PIP3 circuit, which used a much lower time-scaling of 0.040). This determines how quickly time-dependent subsystems (e.g., signaling circuits and diffusion) evolve relative to the stochastic, CPM-driven lattice updates (cell shape changes and movement) when interpreting the simulation in real units. Adjusting these values can create scenarios where cells with even limited feedback (such as WP cells) navigate much more effectively with fewer collisions. However, these specific values were chosen to balance simulation speed and accuracy across all models so that simulation time was reasonable while still allowing some cells to effectively navigate the tracks.

Additionally, to ensure the spatial scale of the reaction-diffusion dynamics is meaningful and not arbitrarily dependent on the specific cell size in pixels used in the CPM, the diffusion terms were scaled by a factor of $1/L^2$, where $L = 2\pi/P_{\text{target}}$ (calculated as $P_{\text{target}} = 2\pi\sqrt{\text{Target Area}/\pi} \approx 132$ in our implementation). This effectively normalizes the physical perimeter of the cell onto a standardized $[0, 2\pi]$ domain, making the reaction-diffusion length scale consistent and mathematically convenient regardless of the simulation's exact spatial resolution. Finally, Rho-induced contraction was necessary for the tension implementation to function properly in Morpheus. Without this contraction, collisions with boundaries were not intense enough to stretch the cell perimeter and trigger the tension effect.

**Table 1.** Morpheus model parameters: values (means and standard deviations, or fixed values) for all model variants (WP, WPI, WPI-PIP3, Rac-Rho, and Rac-Rho-T). WP-variant models and parameter values were previously fit by Algorta et al [30] to data from Town and Weiner [14]. Rates of diffusion for active forms $(u, R, \rho)$ are $0.12/L^2$ and for inactive forms $(v, R_i, \rho_i)$ are $1.0/L^2$, where $L$ is the length scale (see Morpheus implementation specifics). The local Rac inhibitor $h$ diffuses at $0.09/L^2$, and PIP3 $(p)$ diffuses at $0.62/L^2$.

| Symbol | Description | Value / Mean | Std. Dev. |
|---|---|---|---|
| $k_0$, $k_{0,\rho}$ | Basal conversion rate (Rac, Rho) | 0.028 | 0.000765 |
| $k_1$ | Inhibition rate (Rac) | 0.372 | 0.006632 |
| $\alpha_S$ | Stimulus activation coefficient (Rac) | 0.059 | 0.001530 |
| $\alpha_p$ | Stimulus activation coefficient (PIP3) | 0.0068 | 0.000102 |
| $\gamma$, $\gamma_\rho$ | Feedback strength (Rac, Rho) | 1.38 | 0.0214 |
| $\delta$, $\delta_\rho$ | Deactivation rate (Rac, Rho) | 0.721 | 0.006887 |
| $K$ | Activation constant (Rac) | 2.93 | 0.0408 |
| $n$ | Hill coefficient | 2 | – |
| $u, R, \rho$ | Active form of protein | 0.5 | ±0.1 (uniform) |
| $v, R_i, \rho_i$ | Inactive form of protein | 7.31 | ±0.1 (uniform) |
| $h$ | Local Rac inhibitor | 2 | – |
| $p$ | PIP3, signaling lipid | 0.93 | – |
| $T_p$ | Total PIP pool (PIP2 + PIP3) | 3.08 | – |
| $T_0$ | Tension Hill function param (Rac-Rho-T) | 5.0 | – |
| $n_T$ | Tension Hill coefficient (Rac-Rho-T) | 4 | – |
| $\beta$ | Tension strength (Rac-Rho-T) | 3.0 | – |
| Target Perimeter | Target perimeter for tension (Rac-Rho-T) | 172.0 | – |

### Image processing and analysis pipeline

To quantify cell-boundary collisions in our Morpheus CPM simulations, we developed an automated image processing and analysis pipeline, available in the GitHub repository at https://github.com/edwinhuras/morpheus-chemotaxis-analysis. This approach was selected to enable processing similarly structured microscopy data for future comparisons between models and experiments. The repository contains the Morpheus model files, processed collision data, and sample simulation outputs. It also includes the complete processed dataset generated from our analysis pipeline. Because the raw Morpheus output (∼20 GB) exceeds GitHub's storage limits, these files are available from the corresponding author upon request.

The pipeline was developed using Python and standard data science libraries (NumPy, Pandas, OpenCV for image segmentation, and Matplotlib/Seaborn for plotting). AI assistants were utilized to help write the code for data processing and visualization. The pipeline iterates over the simulation outputs and extracts cell trajectories from `celltracks.xml` files when available, otherwise from the simulated images. It processes the PNG frames generated by Morpheus to detect collisions. Specifically, the cell body is identified by black pixel segmentation, and the boundary mask is dilated using a $3 \times 3$ kernel (equivalent to checking the 8-pixel neighborhood). A collision is recorded when any pixel of the cell's perimeter overlaps with the green track boundary pixels. From this, we compute summary metrics including collision percentage (fraction of frames with at least one collision), collision intensity (collision pixels divided by total boundary pixels), and track progress (maximum Y-position divided by domain height).

### Additional figures

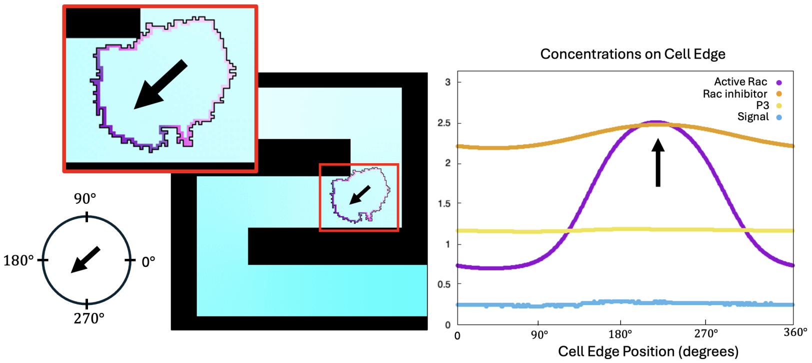


**Fig 13. Visualization of cell polarization and associated protein concentrations during chemotaxis.** A simulated WPI-PIP3 cell in a simple track following a chemical gradient (from white/low to blue/high). On the left, the cell perimeter is shown, colored by the amount of active Rac (highest - purple at the front, lowest - pink at the back) with an arrow pointing in the direction of polarization. The right plot shows distributions of protein concentrations and the detected attractant signal along the cell edge (360° periodic) with respect to the lab frame. Protrusive and contractile behaviors that underlie cell navigation are regulated by membrane protein concentrations evolving according to PDEs, with dynamics simulated in cellular Potts model-based software.

### Movie captions

**S1 Video. Figure 6 Movie (Figure6_Movie.mp4).** Sample cell movements through Track A for each front circuit (WP, WPI, WPI-PIP3) and associated collision metric.

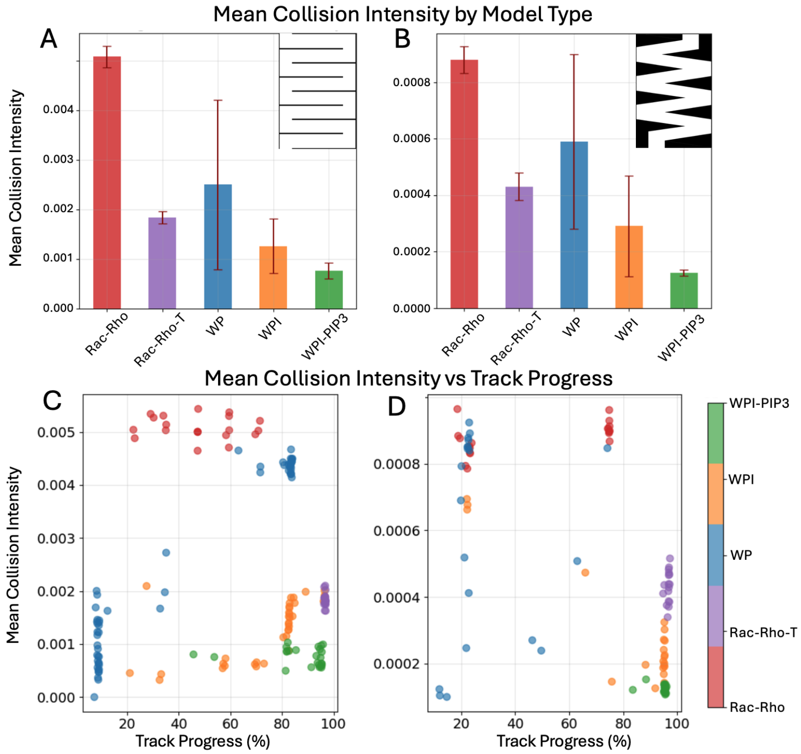


**Fig 14. Summary results for all models.** Rac-Rho and Rac-Rho-T model trials combined with WP variants for comparison. Rac-Rho always polarizes in Track A, unlike WP, although it tends to spend more time stuck in corners and thus makes less overall progress than the successful WP trials. The tension mechanism to Rac-Rho leads to 100% successful track completion with far improved mean collision intensity compared to WP and Rac-Rho.

**S2 Video. Figure 7 Movie (Figure7_Movie.mp4).** As in **S1 Video** but for Track B.

**S3 Video. Figure 10 Movie (Figure10_Movie_truncated.mp4).** Sample cell movements for Rac-Rho front-back circuits (Rac-Rho, Rac-Rho-T) in Track A with WP run for comparison. The Rac-Rho-T model reaches the end of the track quickly and its animation is paused while other models finish.

**S4 Video. Figure 11 Movie (Figure11_Movie_truncated.mp4).** As in **S3 Video** but for Track B.

### Morpheus code

**WP_TrackA.xml, WP_TrackB.xml** *Wave-Pinning (WP) Model of Rac Activation.* This model simulates basic cell polarity via wave-pinning of a single active signalling molecule (Rac). It relies on positive feedback (Rac self-activation) and a conserved total pool of Rac.

**WPI_TrackA.xml, WPI_TrackB.xml** *Wave-Pinning with Inhibitor (WPI) Model.* This model extends the basic wave-pinning (WP) system by adding a local negative feedback loop mediated by a downstream inhibitor of Rac.

**WPI-PIP3_TrackA.xml, WPI-PIP3_TrackB.xml** *Wave-Pinning with Inhibitor and PIP3 (WPI-PIP3) Model.* This model builds upon the WPI system by explicitly incorporating PIP3 dynamics, providing an additional layer of regulation upstream of Rac.

**Rac-Rho_TrackA.xml, Rac-Rho_TrackB.xml** *Rac-Rho Mutual Antagonism Model.* This model explores cell polarization through a mutual antagonism network between Rac (frontness) and Rho (backness), with active Rho promoting cell contraction.

**Rac-Rho-T_TrackA.xml, Rac-Rho-T_TrackB.xml** *Rac-Rho-Tension (Rac-Rho-T) Model.* This model couples a simple tension mechanism, utilizing the cell perimeter as a proxy for membrane tension, to a mutual antagonism network between Rac (frontness) and Rho (backness).
